# The enrichment of breakpoints in late-replicating chromatin provides novel insights into chromoanagenesis mechanisms

**DOI:** 10.1101/2020.07.17.206771

**Authors:** Nicolas Chatron, Giuliana Giannuzzi, Pierre-Antoine Rollat-Farnier, Flavie Diguet, Eleonora Porcu, Tony Yammine, Kevin Uguen, Zohra-Lydia Bellil, Julia Lauer Zillhardt, Arthur Sorlin, Flavie Ader, Alexandra Afenjar, Joris Andrieux, Claire Bardel, Eduardo Calpena, Sandra Chantot-Bastaraud, Patrick Callier, Nora Chelloug, Emilie Chopin, Marie-Pierre Cordier, Christèle Dubourg, Laurence Faivre, Françoise Girard, Solveig Heide, Yvan Herenger, Sylvie Jaillard, Boris Keren, Samantha J. L. Knight, James Lespinasse, Laurence Lohmann, Nathalie Marle, Reza Maroofian, Alice Masurel-Paulet, Michèle Mathieu-Dramard, Corinne Metay, Alistair T. Pagnamenta, Marie-France Portnoï, Fabienne Prieur, Marlène Rio, Jean-Pierre Siffroi, Stéphanie Valence, Jenny C. Taylor, Andrew O. M. Wilkie, Patrick Edery, Alexandre Reymond, Damien Sanlaville, Caroline Schluth-Bolard

## Abstract

The rise of pangenomic molecular assays allowed uncovering complex rearrangements named *chromoanagenesis* that were hypothesized to result from catastrophic shattering events. Constitutional cases have typically been reported individually preventing identification of common features and uncovering the mechanisms at play. We characterized 20 new *chromoanagenesis* and discovered yet undescribed features. While literature differentiates *chromothripsis* and its shattering event repaired through non-homologous end joining from *chromoanasynthesis* born to aberrant replicative processes, we identified shattered chromosomes repaired through a combination of mechanisms. In particular, three samples present with “rearrangement hubs” comprising a fragmented kilobase-long sequence threaded throughout the rearrangement.

To assess the mechanisms at play, we merged our data with those of 20 published constitutional complex chromosomal rearrangement cases. We evaluated if the distribution of their 1032 combined breakpoints was distinctive using bootstrap simulations and found that breakpoints tend to keep away from haplosensitive genes suggesting selective pressure. We then compared their distribution with that of 13,310 and 468 breakpoints of cancer complex chromosomal rearrangements and constitutional simple rearrangement samples, respectively. Both complex rearrangement groups showed breakpoint enrichment in late replicating regions suggesting similar origins for constitutional and cancer cases. Simple rearrangement breakpoints but not complex ones were depleted from lamina-associated domains (LADs), possibly as a consequence of reduced mobility of DNA ends bound to lamina.

The enrichment of breakpoints in late-replicating chromatin for both constitutional and cancer *chromoanagenesis* provides an orthogonal support to the premature chromosome condensation hypothesis that was put forward to explain *chromoanagenesis*.

## Introduction

Since the identification of a supernumerary chromosome 21 in Down syndrome (Lejeune et al. 1959), each technological development (chromosome banding, FISH, cytogenetic microarray, massive parallel sequencing, long-read sequencing) has further exposed the complexity and variability of human chromosomes (Caspersson et al. 1970; Cooper et al. 2011; Sudmant et al. 2015). This “entropy” climaxed with the identification of extremely complex chromosomal rearrangements (CCRs) both in cancer (Stephens et al. 2011) and constitutional samples (Kloosterman et al. 2011) christened *chromothripsis*. These CCRs have their breakpoints clustered to a single chromosome, a chromosome arm or a cytoband and their copy number profile oscillating between two states (1 and 2 copies), in contrast to the classical tumorigenesis scenario. The equal amount of inverted and non-inverted breakpoint-junctions is consistent with a single catastrophic shattering event (*-thripsis* in Greek) followed by random patching through non-homologous end-joining coupled to fragment loss (Korbel and Campbell 2013). The identification of different copy number profiles and alternative replicative repairing mechanisms (Microhomology-mediated breakage induced repair (MMBIR) / Fork Stalling and Template Switching (FoSTeS)) prompted the definition of *chromoanasynthesis* (Liu et al. 2011) and *chromoplexy* involving several chromosomes and only observed in cancer (Baca et al. 2013), with the three phenomena being grouped under the umbrella term *chromoanagenesis* (Holland and Cleveland 2012).

There are two main hypotheses for the underlying mechanisms, mostly deriving from cancer cell observation and cellular models. Firstly, a non-spindle bound lagging chromosome is segregated in a micronucleus (Ganem et al. 2009) where DNA replication is deficient (Crasta et al. 2012; Terzoudi et al. 2015) generating two asynchronous compartments, the micronucleus and main nucleus. This replicating and entrapped chromosome undergoes premature chromosome condensation, which is associated with pulverization of chromatin (Kürten and Obe 1975; Obe and Beek 1975). The second model relies on the dicentric chromosomes (Titen and Golic 2008). Dicentric chromosomes can create bridges between the cytokinesis poles that can only be resolved through multiple DNA breaks (Maciejowski et al. 2015), sometimes with entrapment in a micronucleus (Pampalona et al. 2016). Other hypotheses (incomplete apoptosis (Tubio and Estivill 2011), hyperploidy (Mardin et al. 2015) or mobile elements) have also been formulated.

The observed continuous rather than multimodal distribution of the number of breakpoints from simple to highly complex rearrangements was suggested to mirror a common origin and mechanism (Storchová and Kloosterman 2016). Would this continuum hypothesis be correct, we would expect a similar distribution of breakpoints in simple and complex events. However, beyond low copy repeats known to be elective sites for recurrent non-allelic homologous recombination events (Stankiewicz and Lupski 2002), we are ignorant about possible relationships between genomic context (sequence, genome folding, genome properties) and breakage probability, and potential differences in regard to complexity of the rearrangement.

Here we take advantage of three cohorts of rearrangements – simple, constitutional CCRs and cancer CCRs – to assess and compare the genomic features of breakpoints and provide evidence for the previously hypothesized continuum between simple and complex events. We suggest new elements in the potential mechanisms at play.

## Results

### Chromoanagenesis characterization reveals new molecular features

We recruited 14 unbalanced and 6 balanced *chromoanagenesis* cases. To be included and genome sequenced unbalanced cases had to carry a minimum of 3 non-recurrent non-polymorphic CNVs identified through array-CGH on a single chromosome (potentially chromosome pair) while balanced cases with a minimum of 10 breakpoints were obtained from our translational research study (Schluth-Bolard et al. 2019). Through paired-end short-read genome sequencing, we identified thirteen to hundred breakpoints per rearrangement involving from one to fifteen chromosomes for a total of 682 breakpoints (Table 1 and Figure 1). A detailed description of each case is provided as supplementary data (Supplementary File S1).

**Table 1:** Rearrangements’ characteristics summary. We only inferred the junctions for which a same repairing mechanism was considered at both ends. Full details can be found in Supplementary File S1.

| Individual | No of chromosomes | No of breakpoints | Main chromosome | Copy Number Variant |  | No of resolved junctions | No of inferred junctions | Inferred mechanism |  |  | Inferred mechanism for rearrangement |
| --- | --- | --- | --- | --- | --- | --- | --- | --- | --- | --- | --- |
|  |  |  |  | Gain in Mb (n gain) | Loss in Mb (n loss) |  |  | NHEJ | MMBIR (MMEJ) | Ambiguous |  |
| 1 | 1 | 14 | 3 (100%) | 0 | 0 | 13 | 4 | 1 | 3 | 0 | NA |
| 2 | 1 | 16 | 11 (100%) | 2.25 (2) | 7.15 (4) | 9 | 7 | 3 | 2 | 2 | Chromoanasythesis |
| 3 | 1 | 18 | 6 (100%) | 1.43 (7) | 6.56 (1) | 12 | 9 | 1 | 5 | 3 | Chromoanasythesis |
| 4 | 1 | 18 | 21 (100%) | 5.30 (5) | 11.73 (4) | 9 | 6 | 5 | 1 | 0 | Chromoanasythesis |
| 5 | 1 | 23 | 5 (100%) | 0 | 0 | 23 | 18 | 10 | 5 | 3 | Chromothripsis |
| 6 | 1 | 24 | 20 (100%) | 6.65 (11) | 0 | 14 | 10 | 3 | 3 | 4 | Chromoanasythesis |
| 7 | 1 | 25 | 21 (100%) | 4.54 (11) | 0.31 (1) | 14 | 14 | 3 | 8 | 3 | Chromoanasythesis |
| 8 | 1 | 62 | 7 (100%) | 0.76 (15) | 19.68 (3) | 27 | 27 | 13 | 7 | 7 | Chromoanasythesis |
| 9 | 1 | 68 | 11 (100%) | 8.74 (14) | 4.40 (1) | 52 | 44 | 11 | 23 | 10 | Chromoanasythesis |
| 10 | 2 | 13 | 1 (76,9%) | 0.83 (5) | 0 | 8 | 8 | 8 | 0 | 0 | Chromoanasythesis |
| 11 | 2 | 18 | 2 (94%) | 0 | 2.97 (2) | 15 | 9 | 3 | 6 (1) | 0 | Chromoanasythesis |
| 12 | 2 | 46 | 3 (98%) | 0 | 9.00 (5) | 48 | 29 | 12 | 9 | 8 | Chromoanasythesis |
| 13 | 2 | 56 | 13 (98%) | 35.91 (15) | 15.24 (8) | 33 | 31 | 5 | 24 | 2 | Chromoanasythesis |
| 14 | 3 | 18 | 1 (72%) | 0 | 0 | 16 | 10 | 7 | 0 | 3 | Chromothripsis |
| 15 | 3 | 23 | 2 (91,3%) | 0 | 0 | 18 | 7 | 4 | 3 | 0 | NA |
| 16 | 4 | 14 | 14 (43%) | 0 | 0 | 14 | 10 | 5 | 1 | 4 | Chromothripsis |
| 17 | 4 | 19 | 6 (84%) | 0.25 (1) | 14.97 (6) | 14 | 6 | 3 | 3 | 0 | NA |
| 18 | 5 | 18 | 10 (56%) | 0 | 0 | 16 | 11 | 7 | 1 | 3 | Chromothripsis |
| 19 | 6 | 84 | 4 (68%) | 0 | 6.26 (16) | 71 | 28 | 8 | 12 (2) | 8 | Chromoanasythesis |
| 20 | 15 | 105 | 14 (67%) | 0 | 4.23 (15) | 85 | 54 | 39 | 5 | 10 | Chromoanasythesis |
|  | **Total** | **682** |  |  |  | **511** | **341** | **151** | **121** | **70** |  |

**Figure 1:**
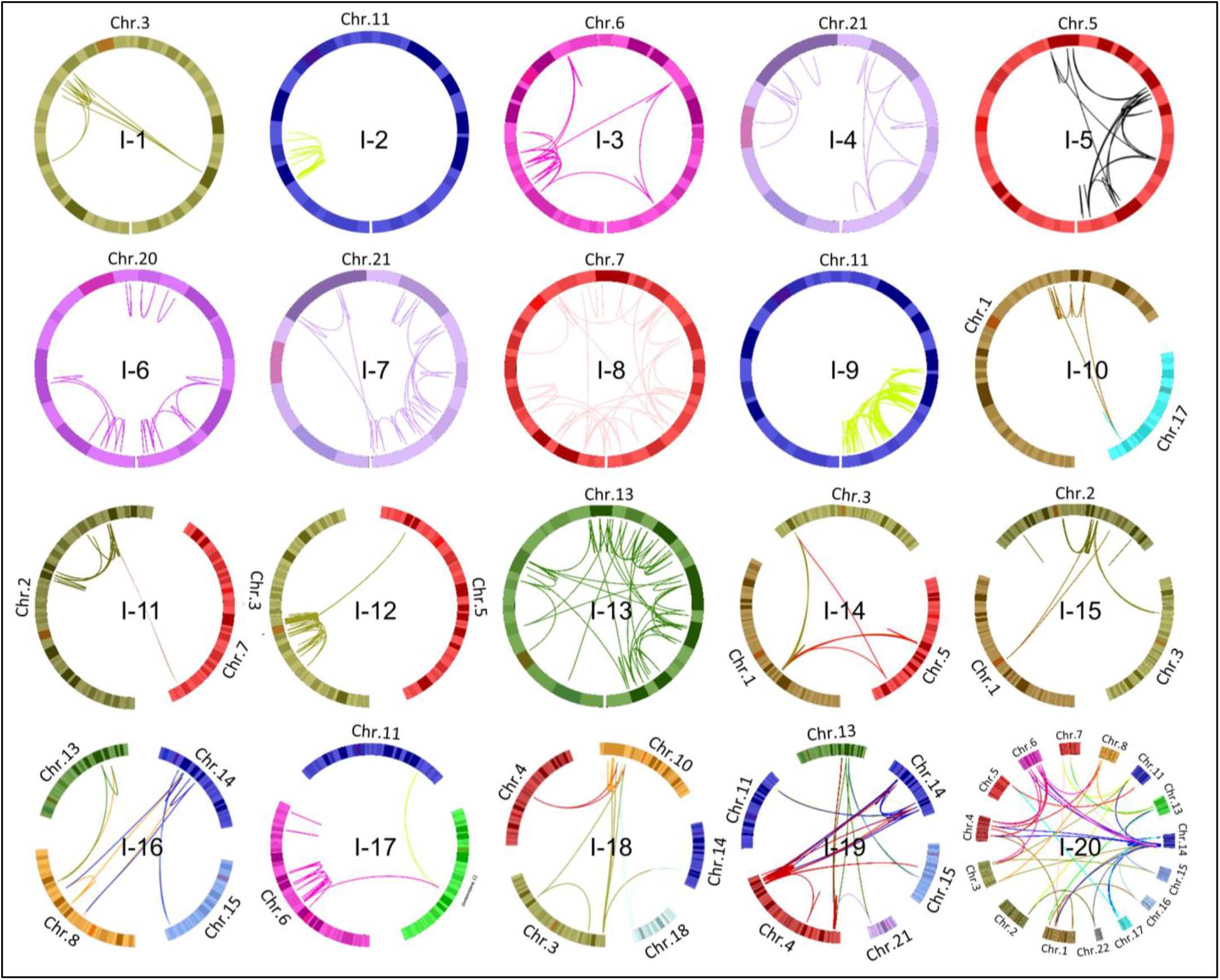
Circos plot visualization of all 20 rearrangements characterized herein. Foci of clustered breakpoints are observed in a majority of individuals.

This characterization effort uncovered noteworthy features. Individual 20 (I-20) carries the most complex *chromoanagenesis* described to date with a single derivative chromosome 14 made of sequences originating from 15 different chromosomes. At several chromosome 14 breakpoints, we observe a “juxtapositions of sequences” from chromosome to chromosome before joining chromosome 14 again until the next “juxtaposition”. Two other single-chromosome cases (I-3 and I-7) also present with copy number gains embedded in junctions but all gains are chained and inserted as a single block. I-10 also shows a single chain of gains inserted as a block. This case is unique as the five gains all derive from chromosome 1 but are inserted in a different chromosome (chr17) close to a paracentric inversion (Figure 2). For these three cases (I-3, I-7 and I-10), the number of junctions not implicating a copy number gain boundary is very limited suggesting that a chromosome shattering is an unlikely cause.

**Figure 2:**
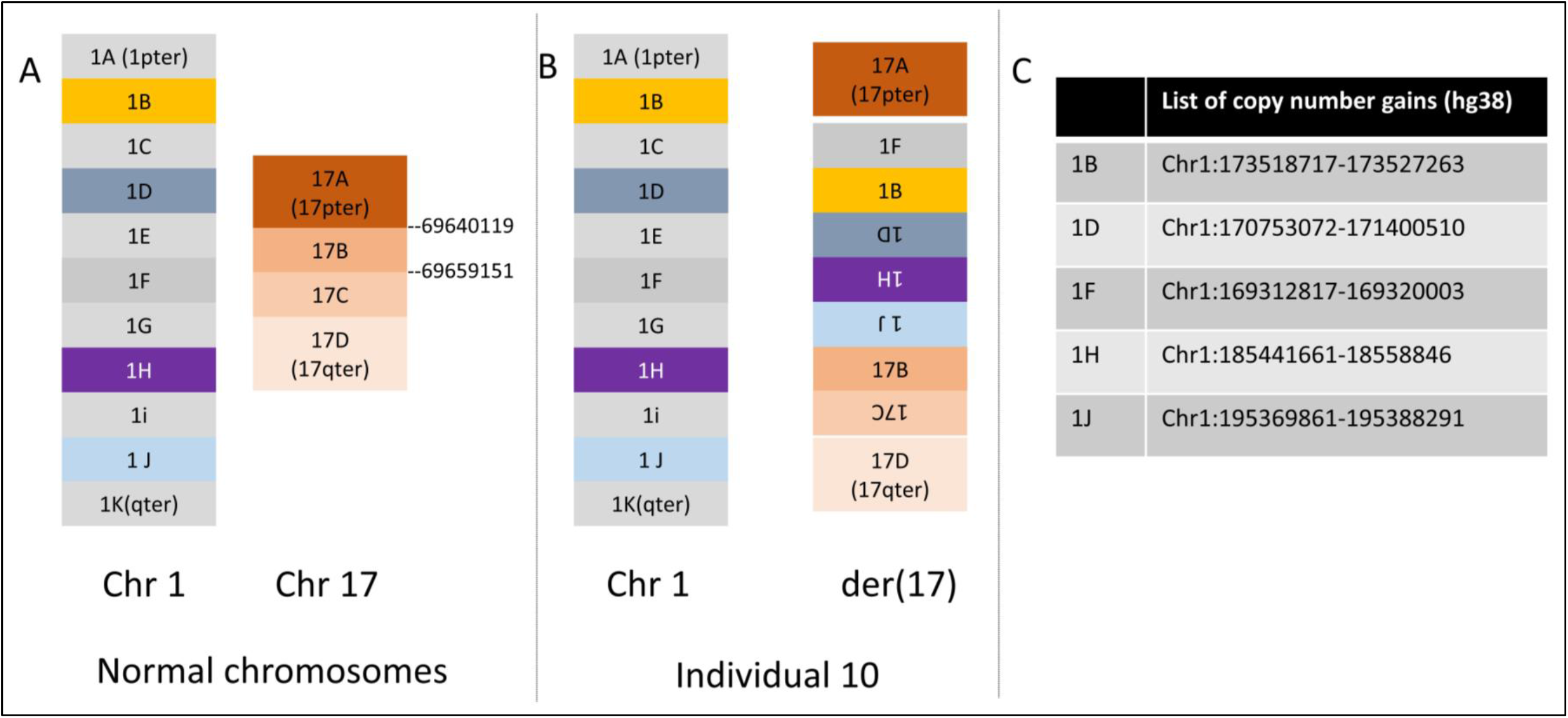
Schematic reconstruction of I-10’s rearrangement: 5 sections of chromosome 1, 1F, 1B, 1D, 1H and 1J (their coordinates are indicated in panel C) are sequelntially inserted in chromosome 17 nearby a 17C paracentric inversion of this chromosome.

Of the remaining rearrangements with multiple chromosomes implicated, I-19 carries the second most complex rearrangement with six chromosomes involved. However, his profile is very different from that of individual 20 as chromosomes 4, 13 and 14 all present with a shattered profile with at least 5 breakpoints each and many junctions connecting pairs of these three chromosomes suggesting that they all had been shattered. Chromosome 11 is only involved in this rearrangement through a 6 kb duplication inserted within a derivative chromosome. I-16 (chromosomes 8 and 14) and I-18 (chromosomes 3 and 10) also harbor more than one shattered chromosome.

Rearrangements of individuals 12, 13 and 15 are a composite of a shattered chromosome and one or two translocations, a situation compatible with a preexisting parental balanced translocation (absent here) or a two-step process during a single gametogenesis.

### Genotype/phenotype correlation

All individuals, but one, of our cohort had an abnormal phenotype. I-7 was identified after a positive first trimester screening for trisomy 21 that revealed an abnormal chromosome 21. Fifteen individuals presented with different degrees of intellectual disability (ID) from mild (I-1, I-18) to severe (I-14). When present ID was associated with different congenital malformations in all individuals but I-6 and I-17. Of the four individuals without ID, two were not assessed formally as they were identified prenatally (I-13) or in the first days of life (I-12). We did not observe any correlation between the phenotype severity and the rearrangement complexity regarding either number of breakpoints, number of chromosomes involved or genomic imbalance. For several individuals the phenotype could be partially explained by the identification of disrupted genes or a position effect. *FOXP1* (Mental retardation with language impairment (MIM 613670)) (I-1), *MEF2C* (Mental retardation, autosomal dominant 20 (MIM 613443)) (I-5 and I-14), *ADNP* (Helsmoortel-van der Aa syndrome (MIM 615873) (I-6), *ZEB2* (Mowat-Wilson syndrome (MIM 235730)) and *COL3A1* (vascular Ehler-Danlos syndrome (MIM 130050)) (I-11), *YY1* (Gabriele-de Vries syndrome (MIM 617557)) (I-20) could partly drive the respective phenotypes. For I-2, 3, 4, 8, 9, 10, 12, 13, 17 and 19 no major gene could be convincingly identified as driving the phenotype but the genomic imbalance was considered large enough to be causative. The striking absence of phenotype for I-7 while she carries a total genomic imbalance of 4.85 Mb on chromosome 21, not affecting the Down Syndrome critical region, could be explained by the insertion of all copy number gains together on the short arm of chromosome 21. Placed in a heterochromatin context the copy number gains could be silenced through a protective position effect.

### Repairing mechanisms are mixed within rearrangements and create “hubs”

We reached nucleotide resolution in both forward and reverse directions for 511 junctions allowing inferring the repair mechanisms through analysis of junction sequence (Supplementary File S1). Using stringent inferring criteria (Online Methods), 162 out of 511 junctions could not be classified. Non-homologous end-joining (NHEJ) was inferred for 151 junctions (30%), while 121 junctions (24%) fitted with our criteria for MMBIR / FoSTeS. Three of these 121 replicative junctions could also be considered as Microhomology Mediated End Joining (MMEJ) as they had no distinctive feature for either mechanism. For 70/341 junctions (14%) we could not distinguish between NHEJ and MMBIR/FoSTeS. Using our criteria to infer subclasses of CCR we classified 13 cases as *chromoanasynthesis* and 4 as *chromothripsis*, leaving 3 rearrangements without clear denomination, mostly due to a limited number of inferable junctions. A single rearrangement in I-10 could be fully inferred with a unique repairing mechanism (NHEJ), whereas all others presented different levels of ascertained mixed signatures and breakpoint distribution was not shown to be significantly different between the two subclasses (data not shown thus questioning the relevance of criteria used for classification and existing definitions.

At sequence resolution, we describe for the first time features that we name “rearrangement hubs”. They consist of short genomic intervals (<1 kb) contacted by more than four junctions each coming from a different locus and using a few dozens of nucleotides of the hub before restarting another rearrangement loop. We identified 5 of these in I-9 (n=2), 12 (n=2) and I-19 (n=1). The most complex “hub” has 12 junctions within a 211 nucleotide span (I-12; Figure 3).

**Figure 3:**
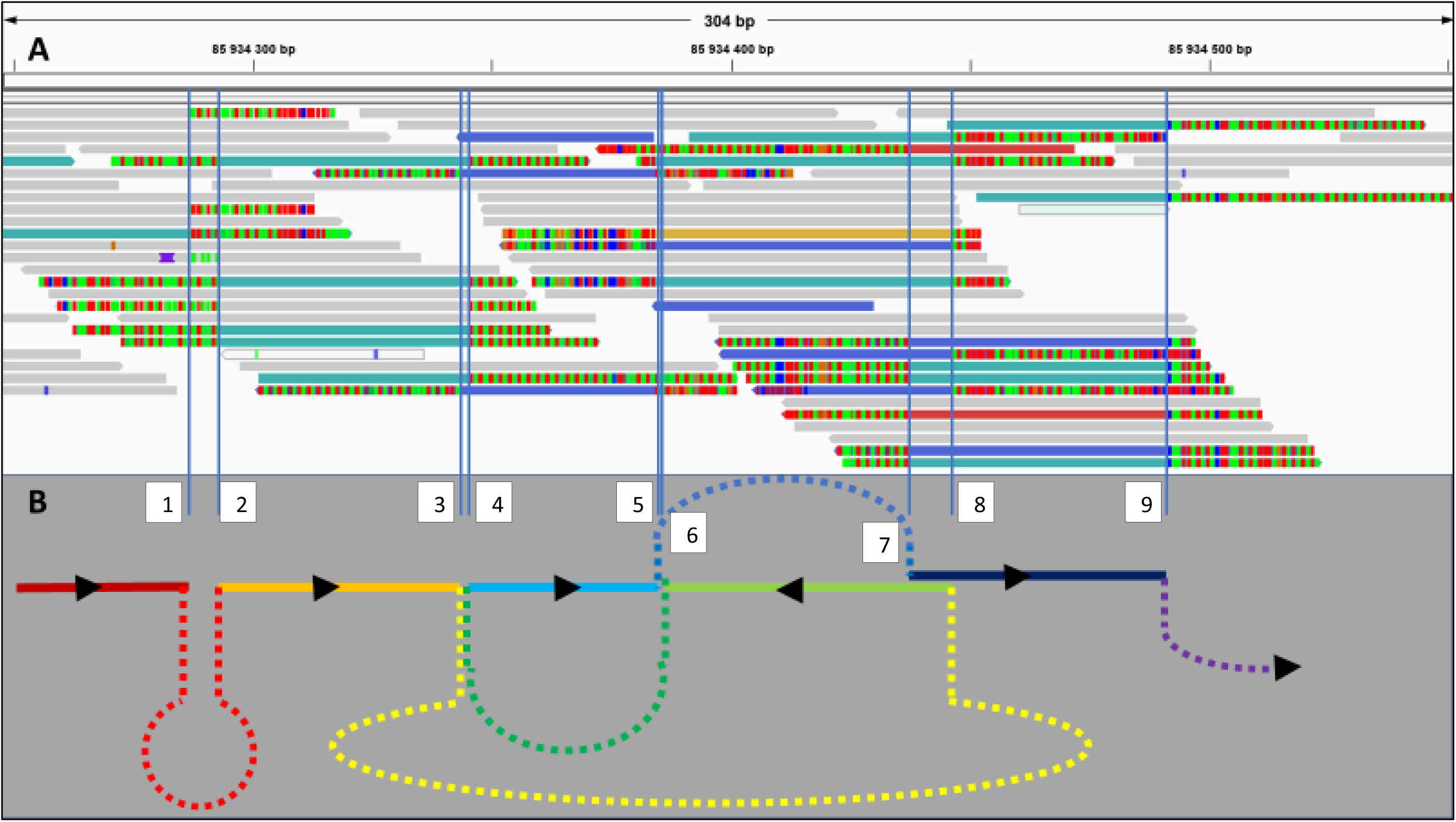
IGV browser (v2.4) view of read alignment of the largest “rearrangement hub” observed in our cohort (I-12) consisting of 9 breakpoints within a segment of 211 nucleotides (A). Reads paired with a mate aligned to a different locus are coloured. Partial schematic reconstruction of the rearrangement with plain lines being sequences of the “hub” and dashed-lines depicting other fragments of the rearrangement. The direction is indicated by black arrowheads (B).

### Chromoanagenesis breakpoint distribution is not random

We then investigated CCR breakpoint distribution to gain insights on potential biological mechanisms. To increase statistical power, we combined 682 breakpoints of our 20 constitutional CCRs with 350 breakpoints of 20 published cases (Yang et al. 2015; Redin et al. 2017). To investigate if the mechanisms involved in constitutional and cancer CCRs are similar we compared the distribution of these 1032 breakpoints to that of 13,310 breakpoints of cancer CCRs deposited in ChromothripsisDB (Yang et al. 2015). We also compared their distribution with that of 468 breakpoints of 234 simple rearrangements described in (Redin et al. 2017) and (Schluth-Bolard et al. 2019) to test the possible continuum from simple to complex rearrangements. We assessed if the breakpoint distribution of these three different groups of rearrangements were enriched within diverse genomic features, such as haplosensitive genes, repeated elements, G-band staining, chromosomal fragile sites, A/B chromatin compartments, lamina associated domains (LADs), topologically associating domains (TADs), replication origins and replication timing. We are aware that these variables are not independant, LADs, for example, are known to be late-replicating gene poor regions (Supplementary Figure S1). Direction of the distribution (enrichment/depletion) were gauged by comparing to two distinct breakpoint simulations (see Online Methods). All annotations are presented in Supplementary Table S1.

First, through univariate analysis, we observed a depletion of both gene-disrupting (Relative Risk Ratio (RRR) = 0.818, Confidence Interval 95% [CI95%] : [0.722-0.927]) and TAD-disrupting (RRR = 0.867 [0.767-0.980]) breakpoints in constitutional CCRs (Figure 4 and Supplementary Table S2). In addition to a massive genomic imbalance, we surmise that the accumulation of disrupted or dysregulated genes would reduce fitness. For simple rearrangements gene-disrupting breakpoints tend to affect haplosensitive genes (higher loss-of-function intolerance (pLI); RRR= 2.267 [1.637-3.137]), probably mirroring the ascertainment bias in recruiting these affected individuals. On the contrary we observed a depletion of high pLI genes in CCRs (RRR= 0.746 [0.585-0.952]), suggesting that their pathologies have an oligogenic basis with multiple breakpoint and CNVs contributing and/or that affecting multiple high pLI genes is incompatible with life. Finally, we observed a significant enrichment in genes and TADs in cancer CCRs (RRR = 1.070 [1.034-1.108] and 1.113 [1.075-1.153] respectively) consistent with a distribution driven by oncogenic forces rather than selective pressure.

**Figure 4:**
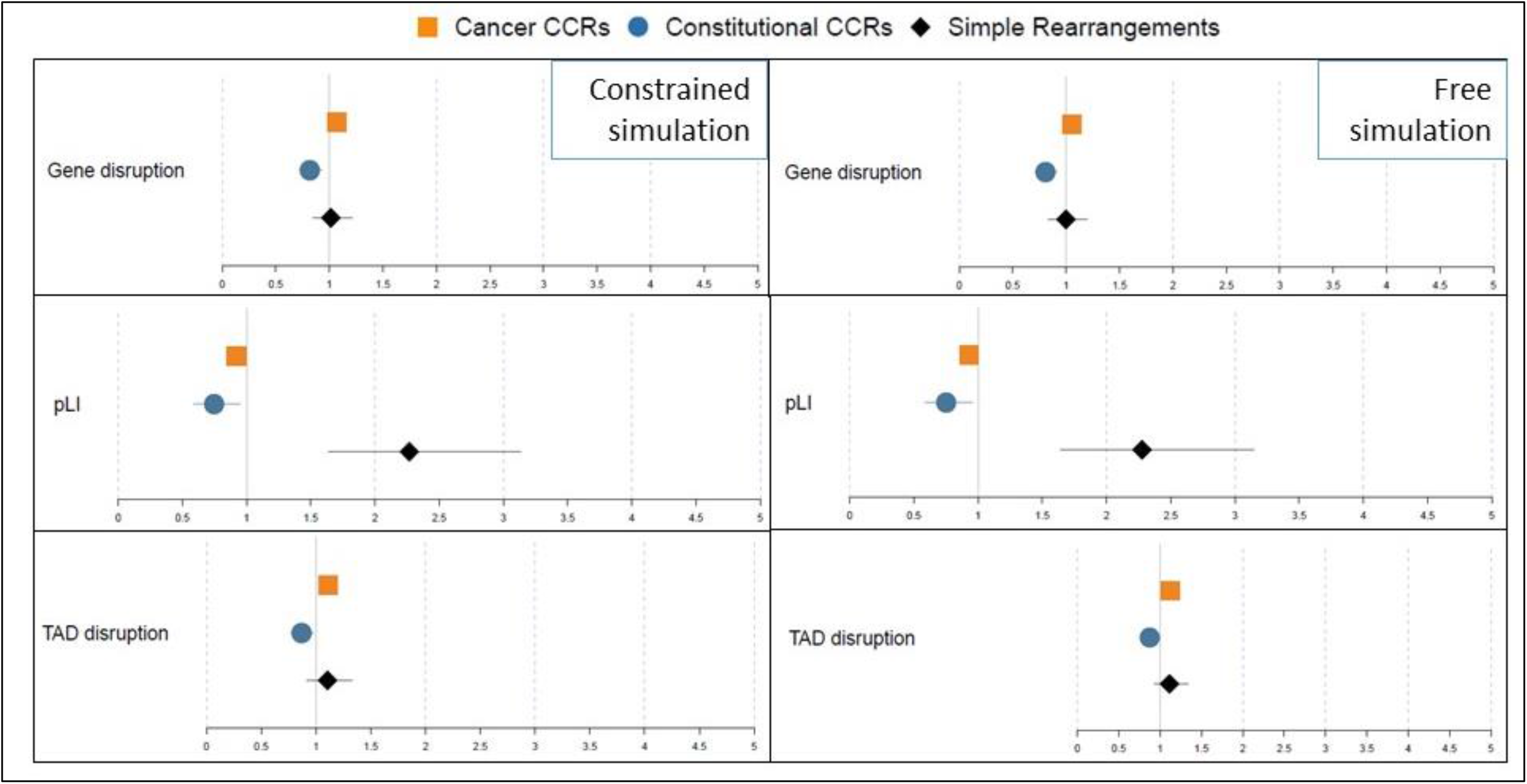
Univariate logistic regression model of breakpoint distribution of simple rearrangements, cancer and constitutional CCRS. These distributions are compared to constrained (left panel) and free simulated distributions (right). Enrichment in genes (top), haplosensitive genes (middle) and TADs (bottom) are shown. Detailed results are presented in table S2. Constitutional complex rearrangements are depleted in gene-disrupting and TAD-disrupting breakpoints as a possible consequence of purifying selection. Simple rearrangements’ breakpoints are enriched in haploinsufficiency intolerant genes possibly reflecting a selection bias towards phenotypically abnormal individuals.

Aside from selective pressure consequences, we identified several genomic features affecting breakpoint distribution. To jointly assess the covariables and confounders (including selective pressure elements) we built a multinomial logistic regression to model the type of rearrangement that originates the breakpoint with the presence/absence of repeated elements, genes, lamina associated domain, and topologically associating domain, the type of chromatin compartment, G-band staining, replication timing status and distance to replication origin (Table 3). In this model, replication timing appears as the unique significant element in the three groups, indeed chromosomal breakpoints are enriched in late-replicating chromatin (RRR = 3.713 [2.583-5.338] for constitutional CCRs; RRR = 1.431 [1.303-1.572] for cancer CCRs; RRR = 2.149 [1.259-3.668] for simple rearrangements) (Figure 5 and Supplementary Table S3).

**Figure 5:**
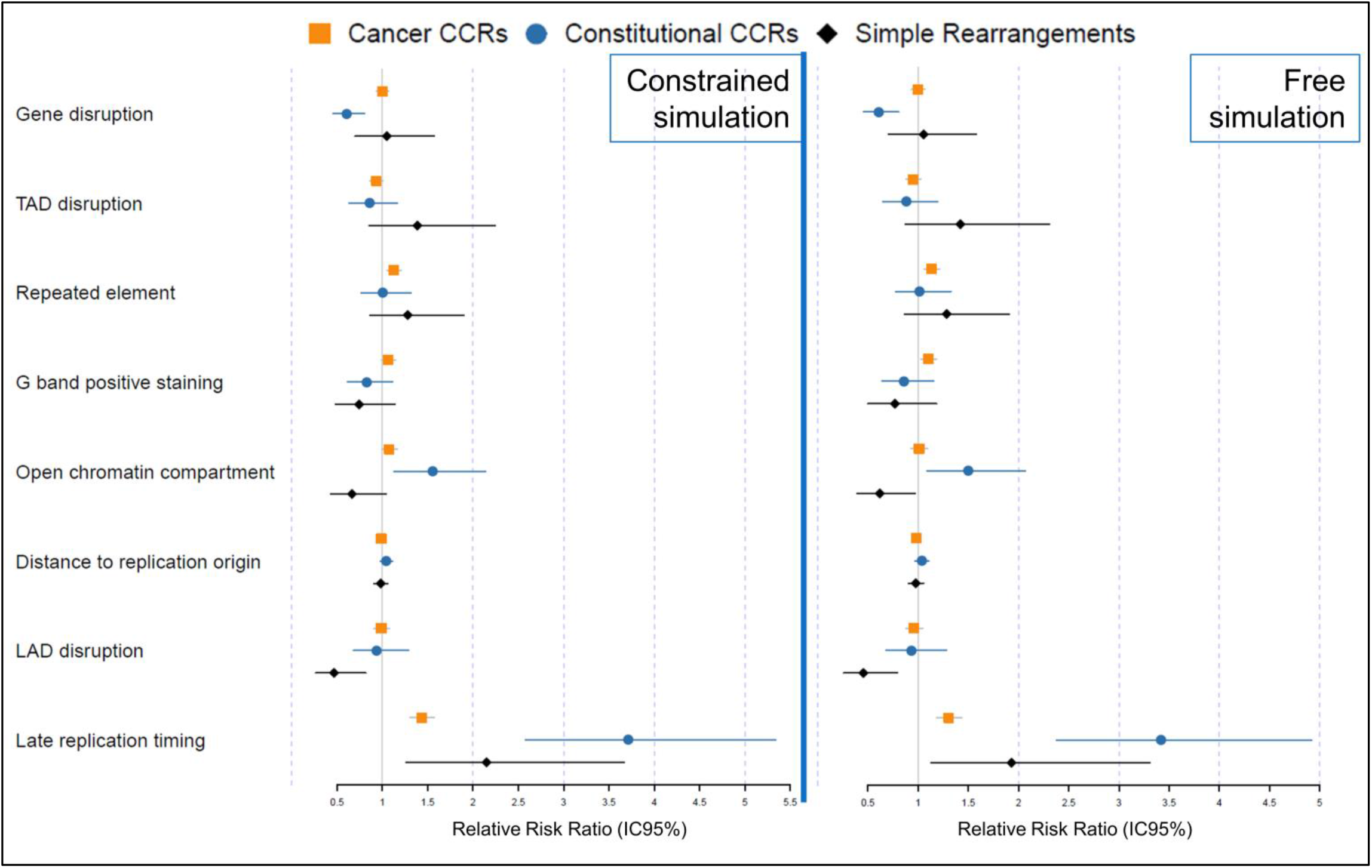
Result of the multinomial logistic regression model of breakpoint distribution of simple rearrangement, cancer and constitutional CCRs. These distributions are compared to constrained (left panel) and free simulated distributions (right) (relative risk ratios and 95% confidence intervals). Distance to replication origin was log-transformed. Late replication timing is the only significant variable for all three groups. The logistic regression model presented is the addition of each covariable that has the lowest Akaike information criterion tested (AIC = 36663,63 and 36558.53 for constrained and free simulation as reference group respectively

For constitutional *chromoanagenesis*, beyond replication timing, depletion in genes (RRR = 0.605 [0.454-0.806]) and enrichment in open chromatin regions (RRR = 1.553 [1.129-2.137]) appear as other significant elements. For cancer rearrangements, none of the analyzed covariables but repeated element disruption was significantly different from simulations so that breakpoint distribution is close to random once adjusted for replication timing.

Finally, while late-replication is known to be a characteristic of LADs, simple rearrangements’ breakpoints appeared depleted within LADs (RRR= 0.466 [0.266-0.816]). Conversely, complex rearrangements’ breakpoints were independent of LADs as their distribution is comparable to breakpoint simulation (RRR= 0.987 [0.905-1.077] and 0.935 [0.679-1.289] for cancer and constitutional cases respectively). Our results suggest that preference towards non-peripheral, i.e. non lamina-associated, chromatin appears as the unique criterion distinguishing the distribution of simple rearrangement breakpoints from that of complex cases that are naive to this feature.

## Discussion

With a total of 20 cases, this is the biggest cohort of constitutional CCRs to date offering new insights on the origins and mechanisms of such rearrangements *in vivo*. Interestingly, of the 13 rearrangements we included based on their abnormal array-CGH result, they all presented additional breakpoints and complexity. The coincidence of three non-recurrent non-polymorphic CNVs identified during array-CGH on a single chromosome is a good indication for an unbalanced CCR requiring genome sequencing for detailed characterization. Of note, paired-end short-read sequencing identifies only 91% of karyotype-visible breakpoints (Redin et al. 2017) suggesting that an extra loop of junctions cannot be ruled out in our “solved” cases.

Our characterization effort challenged several literature definitions. First, we could not disentangle situations where gains are inserted in a single block (I-3, 7 and 10) from replicative repair of a high number of breakpoints in a single chromosome “using” a variety of other chromosomes. This situation questions the definition of *chromoanagenesis* subgroups as it implies a grey zone between *chromothripsis* and *chromoanasynthesis*. In *chromoanasynthesis*, the complexity rather than resulting from chromosome shattering is the consequence of a DNA break. Alternatively, it could stem from a simple replication fork stalling resolved “using” other sequences of the genome through a replicative process (MMBIR/FoSTeS) (Holland and Cleveland 2012; Liu et al. 2011). Thus, what we observe through genome sequencing can either be a DNA breakpoint or a DNA join point. The fact that none of the eight junctions resolved for individual 10 could be inferred as replicative questions our ability to identify a repairing mechanism and differentiate breakpoints from join points. In this “grey zone”, a shattering event seems likely but junction sequences present distinct signs from the classical non-homologous end joining of *chromothripsis* (Kloosterman et al. 2011; Chiang et al. 2012).

Second, we describe a new feature of complex rearrangements that we name “repairing hubs”, where a single locus is “used” several times within the derivative chromosome. We identified five hubs in 3 out of 20 cases; they have no common feature except for G-band positive staining. It will be of particular interest to collect additional observations and further investigate the genomic features shared by these loci. Slamova *et al*. pointed that extremely short sequences can be handled, and here even spread throughout a chromosomal rearrangement, challenging the 50 bp definition of a structural variant (Slamova et al. 2018; Sudmant et al. 2015).

Third, whereas *chromothripsis* has often been used to designate all kinds of *chromoanagenesis* (Redin et al. 2017; Collins et al. 2019; Pellestor 2014; Fukami et al. 2017) our identification of only a few clear cut *chromothripsis* cases in our constitutional cohort suggests that such oversimplification should be avoided.

Apart from variables affected by selective pressure we could not distinguish constitutional *chromoanagenesis* from cancer cases. This suggests that the shattering scenario is comparable for constitutional and tumoral cases. However, while the influence of each variable goes in the same direction for both cancer and constitutional cases, the observed driving forces are still of different magnitude. We hypothesize that this could result from additional complexity delineating possible subgroups. Drier *et al*. have previously shown that cancer rearrangements’ breakpoints have a bimodal distribution, some cancers having breakpoints in early-replicating, highly expressed, GC-rich chromatin while others had their breakpoints preferentially positioned in late-replicating, low transcribed, GC-poor chromatin (Drier et al. 2013). Focusing on CCRs, breakpoints of sarcoma samples are enriched in early-replicating chromatin (Anderson et al. 2018) in contrast to our finding from pooling all cancers together.

By opposition to simple rearrangements, complex rearrangements’ breakpoints appear independent from LADs. First, this is in favor of the micronucleus hypothesis as it was shown that LADs are lost in a majority of micronuclei (Hatch et al. 2013). Second, this draws a demarcation line in the supposed continuum between simple and complex rearrangements emphasizing differences in underlying mechanisms. This LAD depletion is counter-intuitive as, first, the peripheral chromatin was used to be considered as a “body-guard”, absorbing mutagens to the inner chromatin (Hsu 1975) and second, selective pressure should contribute to keep structural variants towards the nuclear periphery as it is both gene-poor and less transcriptionally active.

Simple rearrangements are formed after the mobilization and incorrect pairing of two DNA double strand breaks. This mobilization is restricted to a limited volume so that the pairs formed belong to neighboring chromosomal territories (Soutoglou et al. 2007). Being peripheral a DNA break in a LAD has, by definition, a reduced number of potential partners. DNA breaks in LADs are not mobilized towards nuclear positions more favorable to homologous recombination (Lemaître et al. 2014). Their mobilization capacity might even be reduced by their interaction with lamina, once again reducing their ability to pair with another genomic locus. Overall, we suspect that DNA breaks occurring in LADs are as frequent as elsewhere in the genome but have more chance to be repaired to their native partner than to create a rearrangement compared to “central chromatin”. These events could only be identified at sequence resolution with repairing sequence scars.

In our analysis, late replication timing appears as a key risk factor for *chromoanagenesis* and chromosomal breakage more broadly as it is for other types of genomic variation. It was first associated with important variations in SNP-density across evolution and cancer development with 2-fold and 6-fold enrichment for transitions and transversions mutations respectively (Koren et al. 2012), but profiles are similar in germline and cancer cells (Liu et al. 2013). One possible explanation is that error-prone repairing mechanisms are more active in late replication (Polymerase θ) while early-replicating gene-dense chromatin benefits from high fidelity transcription-coupled nucleotide excision repair (Waters and Walker 2006). Interestingly, polymerase θ had previously been considered to explain the features observed in a *chromoanagenesis* case (Masset et al. 2016). CNV distribution was also reported to be influenced by replication timing, gains and losses being enriched in early and late regions respectively in cancer cells, with breakpoints being part of the same replicating domain. However, to date, there is no robust explanation for such findings (De and Michor 2011). Finally, if partners of balanced rearrangements preferentially replicate at the same time (Ryba et al. 2010) preference towards late replicating regions had never been reported before.

Observation of a breakpoint enrichment in late replicating chromatin supports the “Prematurely Condensed Chromosome” hypothesis for chromoanagenesis. Indeed, our scenario for *chromoanagenesis,* presented in Figure 6, is that after one (or a few) chromosome(s) gets trapped in a micronucleus it can only replicate more slowly than the main nucleus. While entering the next cell division, replication is not completed in the micronucleus and the entrapped material has to start condensation prematurely. This generates a stress on replication forks still present in the physiologically late replicating regions of this delayed chromosome. The forks would either collapse and provoke DNA breakage or stall and start FoSTeS cycles. This would explain the co-occurrence of different repairing mechanisms observed in a same rearrangement. The rearrangement puzzle will be reassembled after reintegration within the main nucleus, opening the possibility for FoSTeS loops to use other chromosomes.

**Figure 6:**
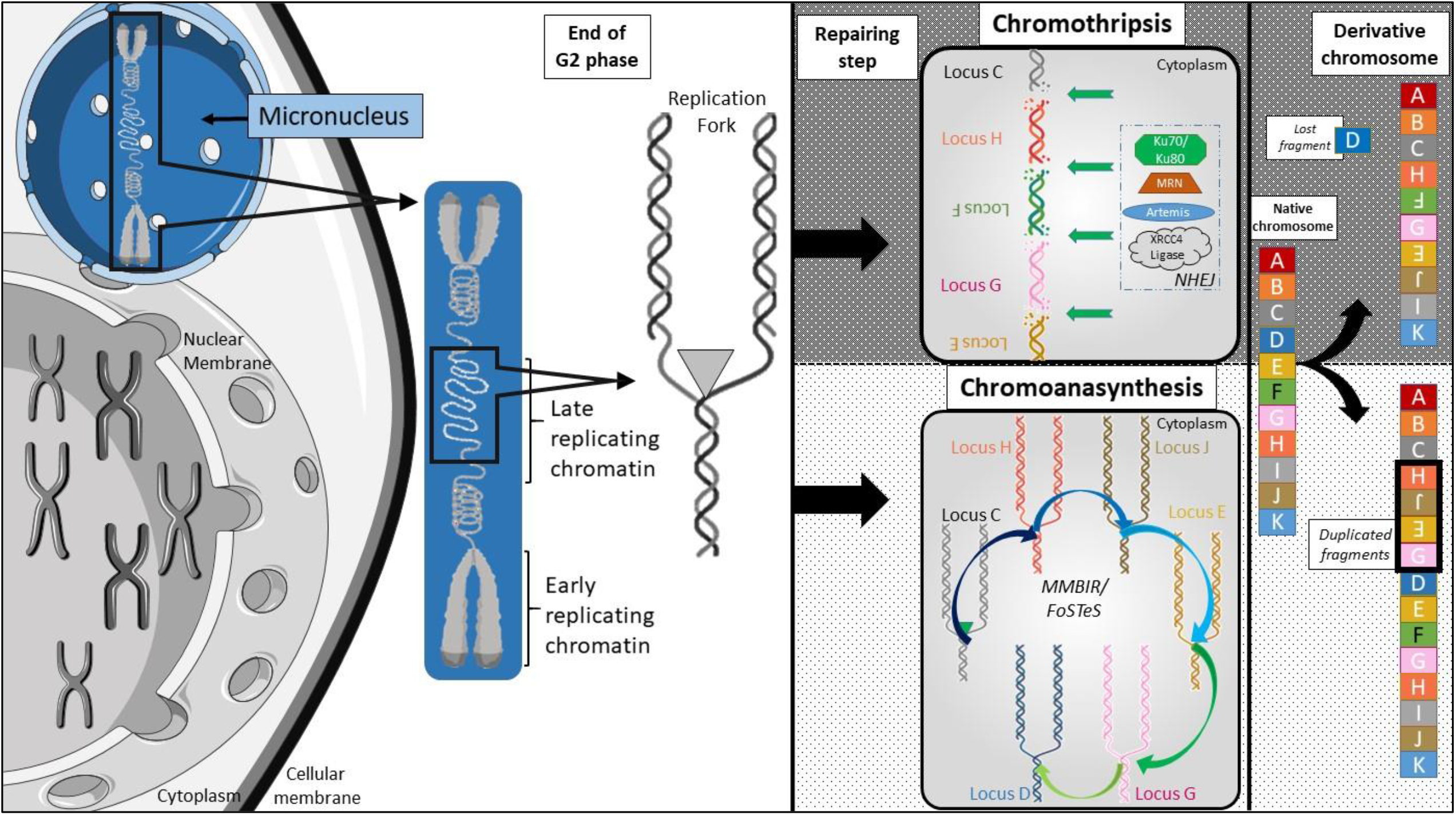
Schematic summary of our scenario to explain chromoanagenesis breakpoint distribution. After missegregation within a micronucleus, DNA replication is delayed. While other chromosomes are ready for cell division, still active replication forks in constitutively late replicating regions are subject to premature condensation and will either collapse (and create DNA double strand breaks) or stall (with possible FoSTeS events) so that both mechanisms can be observed in a single rearrangement.

Finally, the low proportion of rearrangements affecting two chromosomes and the importance of replication-timing is not in favor of a telomere crisis scenario that would involve a dicentric chromosome. Indeed, this second hypothesis can explain clustered breakpoints on a maximum of two chromosomes but requires deficiencies of important cell cycle regulators (*RB, TP53*) that are relatively unlikely during gametogenesis considering constitutional cases. Instead, several rearrangements present a single junction between a shattered chromosome and another one suggesting the preexistence of a balanced translocation. While no constitutional CCR has been described in offspring of simple balanced rearrangement carrier such event could happen in the same gametogenesis and favor entrapment of one chromosome of the tetravalent in a micronucleus.

Overall, we highlight the importance of precise characterization of CCRs as we identified (i) rearrangement “hubs” that had not been described previously and (ii) a “grey zone” between *chromothripsis* and *chromoanasynthesis* challenging actual definitions. By extensive annotation of the largest cohort of constitutional CCRs and use of publicly available data, we identify replication timing as a new element driving chromosomal breakpoint probability. Our *in vivo* dataset supports the implication of premature condensation of genetic material segregated in a micronucleus in *chromoanagenesis*. Finally, we identify a statistically significant depletion of simple rearrangements’ breakpoints within LADs. This strongly suggests that *chromoanagenesi*s and simple rearrangements are not part of a continuous spectrum and opens new perspectives towards chromosomal rearrangement understanding.

## Online Methods

### Individuals

To recruit CCRs we first initiated a French-national call for collaboration to gather all cases analyzed using cytogenetic microarray and having a minimum of 3 non-polymorphic, non-recurrent copy number variants on a single chromosome (potential pair of chromosomes). Individual 10 was recruited to the ethically approved study *Genetic basis of craniofacial malformations* (London - Riverside REC 09/H0706/20) and included after genome sequencing results fitting the entry criterion. In addition, we included all balanced rearrangements characterized in our laboratory as part of our routine activity having more than 10 breakpoints.

All individuals or their parents gave written informed consent for this study, which was conducted with respect to the recommendations of the Helsinki Declaration. Noteworthy, balanced rearrangements had previously been published as part of our clinical validation study (Schluth-Bolard et al. 2019). Individual 20’s description has been recently reported (Ader et al. 2019).

### Genome sequencing

All cases, but case 10, were genome sequenced using our validated approach (Schluth-Bolard et al. 2019). After PCR-free library preparation (Nano, Illumina Inc., San Diego, CA, USA), 2×101 bp paired-end sequencing was performed on a 300 cycles High Output FlowCell for NextSeq500 instrument (Illumina, Inc.). Additionally, samples from individuals I-19, 2, 3, 6, 9, 10 and 19 were 2×151 bp paired-end sequenced on a HiSeq X5 or a HiSeq4000 instrument (Illumina Inc.). Breakpoints were detected using BreakDancer v1.4.5 (Chen et al. 2009) and ERDS v1.1 software was used for CNV calling (Zhu et al. 2012). For all calls, supporting alignment data was systematically inspected using the IGV v2.4 software with soft-clipped reads set as visible (Robinson et al. 2011). At each breakpoint, split-reads were extracted manually to align the soft-clipped sequence using BLAT and obtain fragment junction sequence directly (Kent 2002). For all junctions, the process was repeated in both directions of the junction.

### Breakpoint and junction annotation

Considering extreme complexity, all breakpoints within a kilobase were considered as one for further analyses. We annotated each breakpoint with publicly available datasets using Svagga (https://gitlab.inria.fr/NGS/svagga). RefSeq genes coordinates were extracted from the UCSC Genome Browser; pLi scores were extracted from the gnomAD dataset (Lek et al. 2016). Annotation was also made across lamina associated domains (LADs) (Guelen et al. 2008); topologically associated domains (TADs) (Rao et al. 2014); repeated elements (RepeatMasker); chromosomal cytoband and Giemsa staining (Furey and Haussler 2003); chromosomal common fragile sites (Fungtammasan et al. 2012); replication origins (Massip et al. 2019); A/B compartments (Fortin and Hansen 2015) and replication timing (Dixon et al. 2018). For the latter, only constitutive early and late domains were considered. Both strands of a junction had to be inferred with the same mechanism following the same criteria as in (Schluth-Bolard et al. 2019) to be considered.

Based on existing literature, we established a decisional algorithm to infer the *chromoanagenesis* subgroup for each rearrangement as a whole (Korbel and Campbell 2013; Liu et al. 2011; Baca et al. 2013; Holland and Cleveland 2012). First, rearrangements presenting several copy number gains were classified as *chromoanasynthesis* as the concept of *chromothripsis* does not foresee genomic gains. Second, the rearrangement was considered as *chromothripsis* or *chromoanasynthesis* depending on the dominant repairing mechanism (NHEJ for *chromothripsis;* MMBIR/FoSTeS or clear FoSTeS events with several shifts for *chromoanasynthesis*) with a minimum of 8 inferred junctions to qualify the rearrangement.

### External cases

To increase statistical power we extended our analysis to other complex rearrangements and used ChromothripsisDB (Yang et al. 2015). We downloaded the complete database and kept for analysis samples that had been genome sequenced using paired-end sequencing and for which breakpoint positions were available. We filtered out every call that was (i) not on chromosomes involved in the rearrangement as described by initial authors; (ii) of uncertain position as they had ‘start’ and ‘stop’ positions distant more than 1 kb. We finally extended our cohort of constitutional CCRs with a few more recent constitutional cases (Redin et al. 2017) absent from ChromothripsisDB for a total of 20 external cases (350 breakpoints). Before pooling all constitutional CCRs, we confirmed that our dataset was similar to published cases when considering breakpoint distribution. Compared to simulation data, both distributions were either not significant or in the same direction considering individual genomic features (Supplementary Figure S2).

Constitutional simple rearrangements published in (Schluth-Bolard et al. 2019) and (Redin et al. 2017) were pooled in a third group distinct from (a) constitutional and (b) cancer CCRs.

### Breakpoint simulation

We performed *in silico* breakpoint simulation to create a null hypothesis of breakpoint distribution. We used R software for two different Monte Carlo simulations. In both cases, a random number of breakpoints between 10 (minimal number of breakpoints we used to consider a *chromoanagenesis*) and 1000 were picked for 1000 simulations. In the first “free” simulation, each breakpoint could occur in every possible position of the genome. In the “constrained” model, to mimic the observed “geographical clustering” of breakpoints, a random percentage between 50 and 100% of breakpoints had to occur in the same chromosome while the remaining ones were free. As we know that some genomic regions are poorly analyzed for structural variants, we used the “DangerTrack” bed file (Dolgalev et al. 2017) to exclude all breakpoints simulated in these regions and be comparable with the observed dataset. A single breakpoint of our in-house dataset overlaps the DangerTrack. Overall, 399,346 and 399,915 breakpoints were simulated and used for analyses for the “free” and “constrained” models respectively.

### Statistical analysis

All statistical analyses were performed using R software v.3.5.2. Univariate analysis of each predictor and the logistic regression model building was performed using the “multinom” function of the “nnet” R package (Ripley and Venables 2016) as the Y outcome is multinomial. The Wald test was used for p-value calculation. Considering the number of variables and hypotheses tested we considered p < 0.01 for statistical significance.

## Data access

All sequencing data generated were obtained from patients in a diagnostic setting and can be shared individually upon reasonable request for patients that had consented data sharing.

## Acknowledgments

The authors would like to thank members of the AchroPuce network for their thorough collaboration. NC acknowledges « Fondation Thérèse et René Planiol », Réseau Régional de Rééducation et de Réadaptation Pédiatrique en Rhône Alpes, IDEX Lyon and Hospices Civils de Lyon. GG is recipient of a Pro-Women Scholarship from the Faculty of Biology and Medicine, University of Lausanne. This work was supported by grants from the Swiss National Science Foundation (31003A_182632) and Horizon2020 Twinning project ePerMed (692145) to AR, NHR Oxford Biomedical Research Centre based at Oxford University Hospitals NHS Trust and University of Oxford to AOMW, the Fondation Maladies Rares (WGS20140103) to DS, from the French Ministry of Health (DGOS) and the French National Agency for Research (ANR) (PRTS 2013: PRTSN1300001N) to CSB. Patient 10’s whole genome sequencing was funded by independent research commissioned by the Health Innovation Challenge Fund (JCT), a parallel funding partnership between Wellcome (WT 100127) and the UK Department of Health (R6-388). The views expressed in this publication are those of the authors and not necessarily those of Wellcome, NHS, the NIHR or the Department of Health.

## Author Contributions

Nicolas Chatron, Pierre-Antoine Rollat Farnier, Flavie Diguet, Tony Yammine, Kevin Uguen, Zohra-Lydia Bellil, Julia Lauer Zillhardt, Arthur Sorlin performed the experiments and analyzed the data. Nicolas Chatron, Giuliana Giannuzzi, Claire Bardel, and Eleonora Porcu performed statistical analyses. Flavie Ader, Alexandra Afenjar, Joris Andrieux, Eduardo Calpena, Sandra-Chantot-Bastaraud, Patrick Callier, Nora Chelloug, Emilie Chopin, Marie-Pierre Cordier, Christèle Dubourg, Laurence Faivre, Françoise Girard, Solveig Heide, Yvan Herenger, Sylvie Jaillard, Boris Keren, Samantha J. L. Knight, James Lespinasse, Laurence Lohmann, Nathalie Marle, Reza Maroofian, Alice Masurel-Paulet, Michèle Mathieu-Dramard, Corinne Metay, Alistair T. Pagnamenta, Marie-France Portnoï, Fabienne Prieur, Marlène Rio, Jean-Pierre Siffroi, Stéphanie Valence, Jenny C. Taylor, Andrew O. M. Wilkie, Patrick Edery enrolled patients in the study, collected the clinical data, and analyzed the data. N. Chatron, Giuliana Giannuzzi, Caroline Schluth-Bolard and Alexandre Reymond drafted the manuscript. Damien Sanlaville, Andrew O. M. Wilkie, helped in revising and editing the manuscript. All the authors approved the manuscript submission.

---

**Supplementary Table S1:** List of annotated breakpoints detected in our cohort of complex chromosomal rearrangements (CCRs), in external cases of CCRs (cancer and constitutional), simple rearrangements and simulations. For replication timing, CE, CL and S stand for constitutive early, constitutive late and switching respectively. Only the two first ones were used for statistical analyses.

**Supplementary File S1:** Individual phenotypic and cytogenetic results before genome sequencing. List of junctions resolved in both forward and reverse orientations with the same repairing mechanism obtained from genome sequencing data with inferred repairing mechanism.

**Supplementary Table S2:**
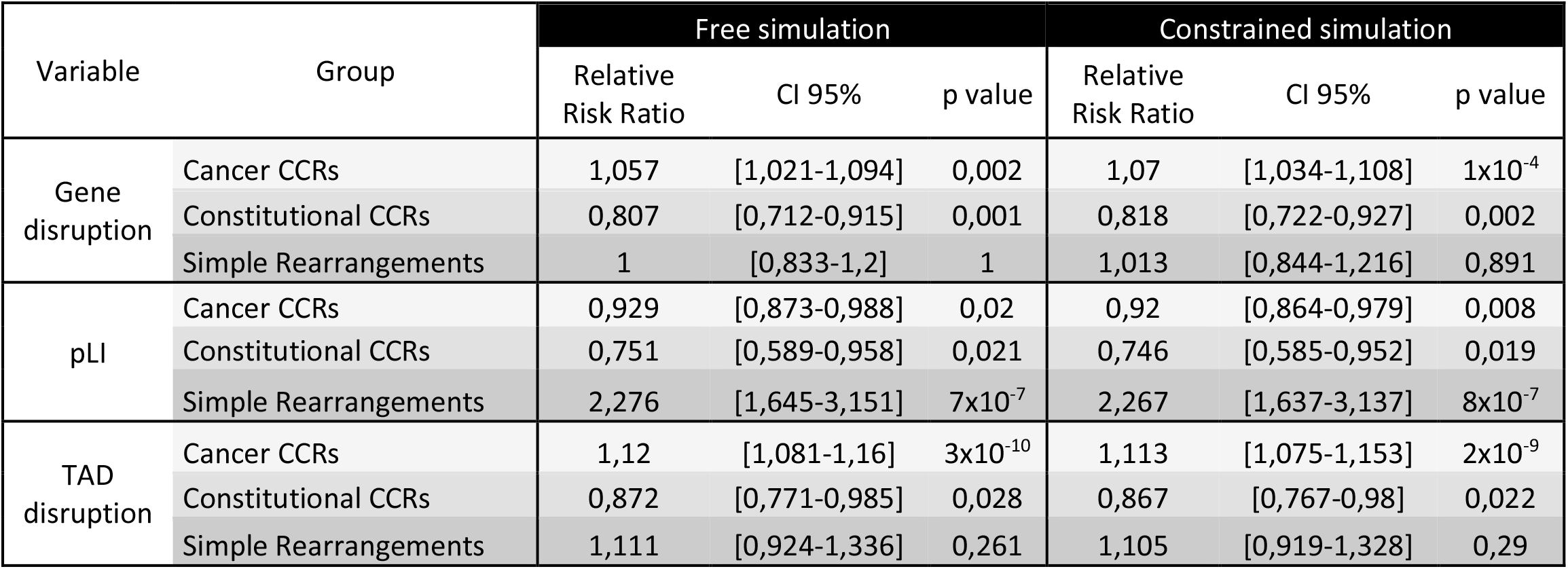
Univariate analysis of breakpoint distribution across genomic features related to gene dysregulation. While constitutional CCRs affect a reduced proportion of genes and TADs compared to simulations, simple rearrangements are more likely to affect haplosensitive genes.

**Supplementary Table S3:** Multivariate analysis of breakpoint distribution across different types of chromosomal rearrangements. The logistic regression model presented is the addition of each covariable that has the lowest Akaike information criterion tested (AIC = 36663,63 and 36558.53 for constrained and free simulation as reference group respectively. Late replication-timing appears as the only significant variable for both cancer and constitutional CCRs. Simple rearrangements diverge from simulation with a depletion in LADs.

| Variable | Group | Free simulation |  |  | Constrained simulation |  |  |
| --- | --- | --- | --- | --- | --- | --- | --- |
|  |  | Relative Risk Ratio | CI 95% | p,value | Relative Risk Ratio | CI 95% | p,value |
| Gene disruption | Cancer CCRs | 0,998 | [0,931-1,069] | 0,955 | 0,998 | [0,931-1,069] | 0,949 |
|  | Constitutional CCRs | 0,606 | [0,455-0,808] | 0,001 | 0,605 | [0,454-0,806] | 0,001 |
|  | Simple Rearrangements | 1,055 | [0,704-1,581] | 0,795 | 1,049 | [0,7-1,572] | 0,818 |
| TAD disruption | Cancer CCRs | 0,950 | [0,878-1,027] | 0,195 | 0,933 | [0,863-1,009] | 0,082 |
|  | Constitutional CCRs | 0,882 | [0,652-1,193] | 0,417 | 0,860 | [0,636-1,162] | 0,326 |
|  | Simple Rearrangements | 1,421 | [0,877-2,303] | 0,154 | 1,386 | [0,856-2,242] | 0,184 |
| Repeated element | Cancer CCRs | 1,135 | [1,061-1,214] | $2 \times 10^{-4}$ | 1,126 | [1,052-1,204] | 0,001 |
|  | Constitutional CCRs | 1,014 | [0,775-1,327] | 0,919 | 1,004 | [0,767-1,314] | 0,977 |
|  | Simple Rearrangements | 1,285 | [0,865-1,908] | 0,214 | 1,280 | [0,862-1,901] | 0,222 |
| G band positive staining | Cancer CCRs | 1,103 | [1,026-1,186] | 0,008 | 1,064 | [0,99-1,143] | 0,094 |
|  | Constitutional CCRs | 0,860 | [0,639-1,156] | 0,317 | 0,826 | [0,614-1,111] | 0,207 |
|  | Simple Rearrangements | 0,769 | [0,5-1,183] | 0,232 | 0,743 | [0,483-1,142] | 0,176 |
| Open chromatin compartment | Cancer CCRs | 1,009 | [0,931-1,094] | 0,824 | 1,074 | [0,992-1,164] | 0,078 |
|  | Constitutional CCRs | 1,499 | [1,087-2,066] | 0,014 | 1,553 | [1,129-2,137] | 0,007 |
|  | Simple Rearrangements | 0,618 | [0,394-0,969] | 0,036 | 0,665 | [0,426-1,037] | 0,072 |
| Distance to replication origin | Cancer CCRs | 0,982 | [0,969-0,995] | 0,008 | 0,988 | [0,975-1,001] | 0,075 |
|  | Constitutional CCRs | 1,036 | [0,971-1,105] | 0,280 | 1,042 | [0,976-1,111] | 0,218 |
|  | Simple Rearrangements | 0,977 | [0,904-1,055] | 0,547 | 0,982 | [0,909-1,061] | 0,647 |
| LAD disruption | Cancer CCRs | 0,960 | [0,88-1,046] | 0,352 | 0,987 | [0,905-1,077] | 0,773 |
|  | Constitutional CCRs | 0,932 | [0,678-1,281] | 0,666 | 0,935 | [0,679-1,289] | 0,683 |
|  | Simple Rearrangements | 0,454 | [0,26-0,793] | 0,006 | 0,466 | [0,266-0,816] | 0,008 |
| Late replication timing | Cancer CCRs | 1,305 | [1,187-1,434] | $3 \times 10^{-8}$ | 1,431 | [1,303-1,572] | $7 \times 10^{-14}$ |
| | Constitutional CCRs | 3,419 | [2,376-4,92] | $3 \times 10^{-11}$ | 3,713 | [2,583-5,338] | $1 \times 10^{-12}$ |
|  | Simple Rearrangements | 1,930 | [1,127-3,306] | 0,017 | 2,149 | [1,259-3,668] | 0,005 |

**Supplementary Figure S1:**
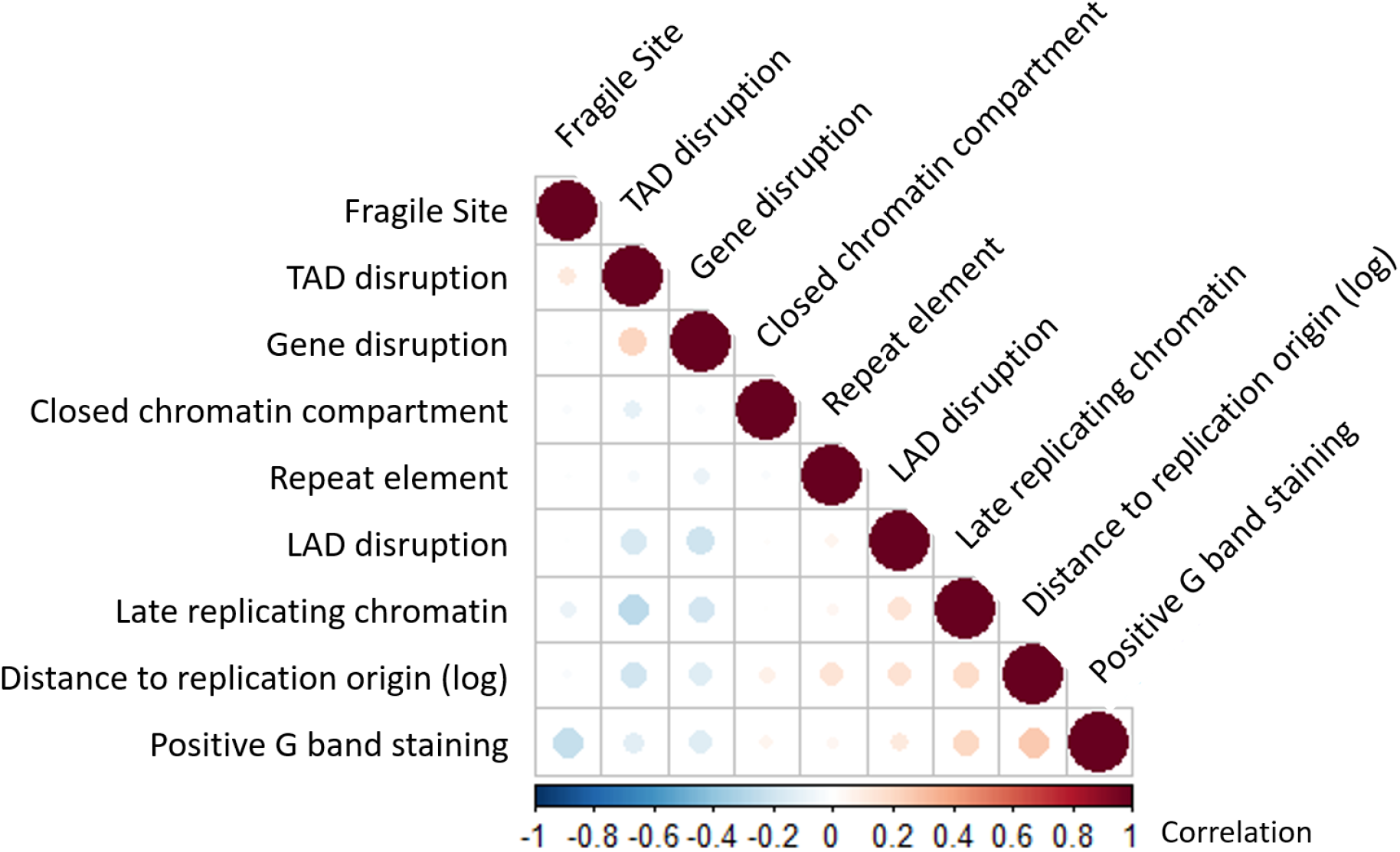
Correlation matrix (Pearson) of the analyzed variables based on our dataset of constitutional CCRs.

**Supplementary Figure S2:**
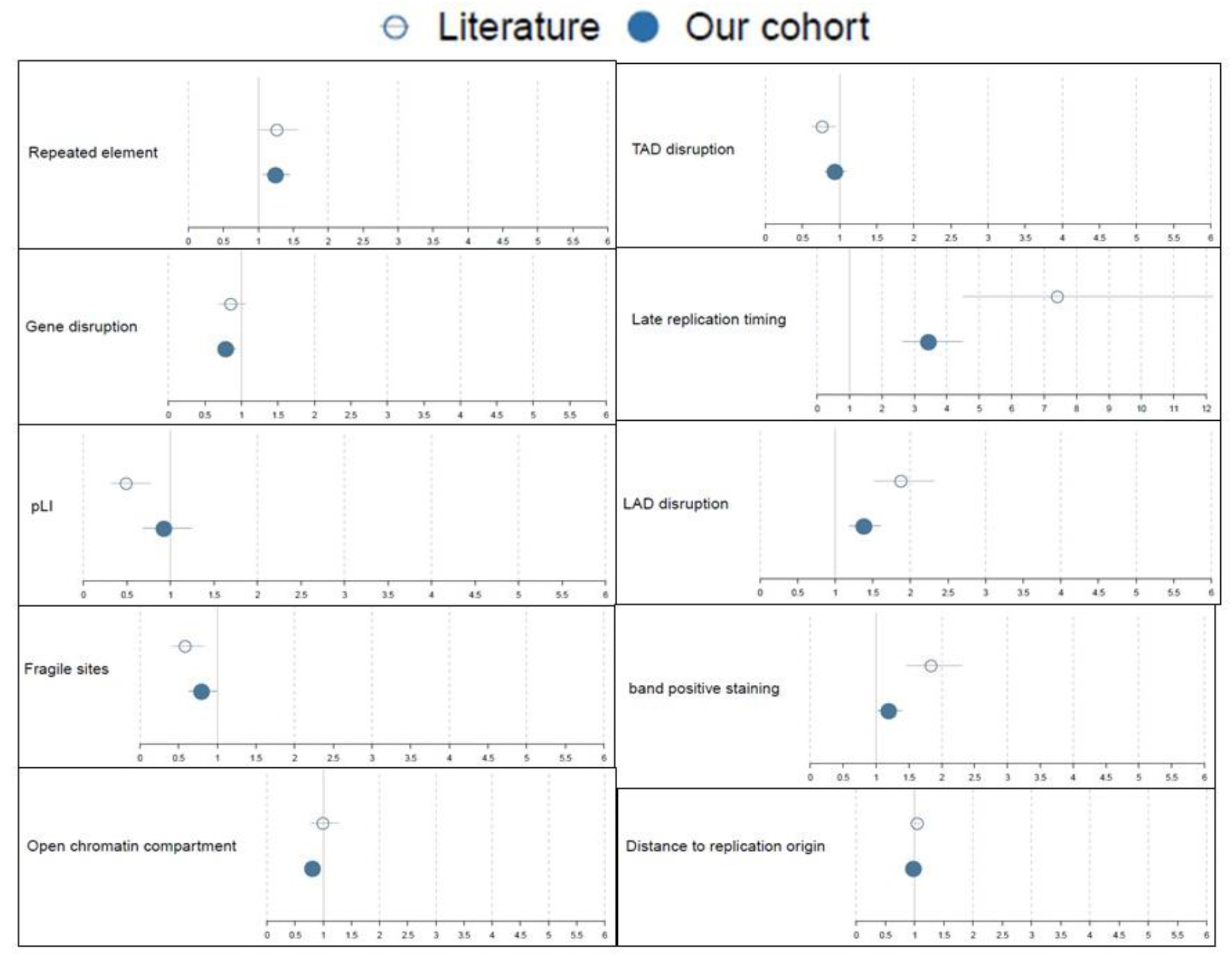
Univariate analysis of breakpoint distribution for our cohort (n=682 breakpoints) and constitutional cases from the literature (n=350 breakpoints) using our “free simulation as a reference group. Relative risk ratios and 95 % confidence intervals systematically overlap and/or are shifted in the same direction so that the two groups can be merged for further analyses.

